# Fetal-intrinsic antiviral mechanisms emerge over the course of gestation

**DOI:** 10.64898/2026.09.22.753282

**Authors:** Drake T. Philip, Madeline S. Merlino, Elizabeth A. Kennedy, Kristen Walsh, Elizabeth Ramos, Gretchen S. Fazakerly, Christopher A. Hunter, Kellie A. Jurado

## Abstract

Congenital viral infections have variable effects on pregnancy outcomes with implications for maternal and fetal health. However, the maternal and fetal immune mechanisms that emerge over the course of gestation to determine protective or pathological outcomes remain poorly understood. Here, we use the emerging congenital pathogen Oropouche virus (OROV) to examine gestational stage-dependent differences in maternal and fetal outcomes in a mouse model of congenital infection. Pregnant mice (dams) infected during early gestation resist severe OROV disease, whereas mid-gestation-infected dams succumb to infection. In contrast, fetal pathology is substantial following early gestation infection but limited following infection during mid-gestation, revealing discordant maternal and fetal susceptibility across gestation. Mid-gestation fetal tissues effectively restricted vertical transmission compared to early gestation fetal tissues, corresponding with reduced fetal pathology. Moreover, both placental and fetal tissue cleared OROV RNA over the course of infection, independent of gestational stage, and failure to clear viral RNA was associated with severe fetal pathology. Spatial analysis of early gestation implantation sites further revealed distinct regional susceptibility to OROV infection across the maternal-fetal interface. We identified potential instances of placental-independent vertical transmission via direct fetal contact with highly infected regions of the contralateral maternal uterus. Finally, we uncovered an unexpected mechanism by which type I interferon signaling contributes to inter-fetal immune crosstalk to restrict both OROV vertical transmission and pathology. Together, these findings establish the fetus as an active participant in antiviral defense and reveal previously unrecognized mechanisms by which fetal-intrinsic antiviral immune responses limit congenital viral infection and disease.

## INTRODUCTION

Immunity during pregnancy is finely balanced, protecting both mother and developing fetus from external threats while maintaining immunological tolerance of the fetus^1^. The maternal and fetal immune systems are highly dynamic and undergo coordinated changes throughout gestation to support fetal development and a healthy pregnancy^1^. Viral infections during pregnancy can disrupt this balance, leading to severe consequences for both the parent and developing fetus^2^. However, our understanding of the mechanisms by which maternal and fetal immune systems sense, control, and clear infection over the entire course of pregnancy remains incomplete. Antiviral immune responses during pregnancy can have distinct outcomes depending on gestational stage^3,4^, underscoring the importance of examining antiviral immunity across the course of pregnancy. For example, although the antiviral cytokine interferon lambda (IFN-λ) is constitutively produced throughout gestation by the fetal placenta^3,5^, its effects are temporally distinct, contributing to maternal-mediated fetal pathology during infection early in gestation while driving maternal-mediated protection from congenital infection as gestation progresses^3^. However, this work, as well as others, primarily focuses upon maternal immunity and its effects on fetal outcomes, leaving fetal-intrinsic immunity largely unexplored.

The placenta is a fetal-derived organ that plays a central role in coordinating antiviral defense during pregnancy, with placental-intrinsic antiviral mechanisms best characterized *in vitro/ex vivo* using placental explant and organoid systems^6^. For example, both IFN-λ and interleukin-27 (IL-27) can restrict congenital viral infection in trophoblast organoids^7^. More recently, trophoblast organoids were used to identify DUX4 as a constitutively expressed trophoblast factor that restricts placental herpesvirus infection^8^. Despite these advances, our mechanistic understanding of fetal antiviral immunity during congenital viral infection remains limited, in part due to the challenging nature of mouse pregnancy models and genetic systems. Recent studies have begun to reveal distinct roles for fetal-intrinsic antiviral signaling during congenital infection. During congenital Zika virus (ZIKV) infection, fetal mitochondrial antiviral signaling protein (MAVS) signaling was shown to have region-specific functions across the placenta^9^. While the fetal placenta was highly dependent on MAVS to limit fetal infection and pathology, maternal immunity in the decidua was largely independent of MAVS^9^. In contrast, signaling via the fetal type I IFN receptor (IFNAR) can be detrimental to fetal health during ZIKV infection, with IFNAR-sufficient fetuses exhibiting substantially greater rates of resorbed fetuses, aggregate of dying fetal and placental tissues, than those that lack IFNAR signaling^10^. Notably, these studies examined infection at distinct gestational timepoints, raising the possibility that gestational stage is an important determinant of whether fetal antiviral immune responses are protective or pathogenic during viral infection.

Oropouche virus (OROV) is a re-emerging *Orthobunyavirus* that has historically circulated within the Amazon basin^11^. However, as of 2024, OROV spread to non-endemic regions causing outbreaks throughout South America and the Caribbean^11,12^. Amidst these outbreaks, increasing evidence has linked OROV infection during pregnancy to adverse outcomes including stillbirth, miscarriage, and microcephaly^12–14^. Further OROV RNA was detected across multiple tissues from a stillborn fetus, providing evidence consistent with vertical transmission^13^. Our group and others have experimentally established that OROV infection can induce fetal pathology in pregnant mice^15–18^, yet the manifestations and underlying biology of congenital OROV infection appear strikingly distinct from those observed during congenital ZIKV infection^15,18^. Therefore, investigating OROV infection at different gestational time points of pregnancy provides an opportunity to understand disease associated with a reemerging pathogen while serving as a new model to uncover fundamental mechanisms of maternal and fetal antiviral immunity.

Here, using a recently established mouse model of congenital OROV infection, we uncover discordant pathologic outcomes between mother and fetus that are dependent upon gestational stage. By defining the kinetics of fetal OROV infection over the course of gestation, we are the first to demonstrate productive vertical transmission to intact fetuses. Further, we identify gestational stage as a key determinant of fetal susceptibility to infection and pathology, independent of maternal disease severity. Remarkably, we found the fetus clears OROV RNA over the course of infection independent of gestational stage, revealing an intrinsic capacity to control viral infection that has not been described in other congenital viral infection models. Using immunofluorescence, we identify regional control of OROV infection throughout the uterus, placenta, and fetus, suggesting multiple coordinated pathways that contribute to antiviral immunity across the maternal-fetal interface. Finally, we find a protective effect for inter-fetal immune crosstalk in controlling OROV vertical transmission and pathology. Overall, our study highlights the critical contributions of the fetus to controlling congenital viral infection and limiting adverse outcomes, opening up new avenues to define the mechanisms that coordinate fetal antiviral immunity and protect fetal health.

## RESULTS

### Gestational stage shapes discordant maternal and fetal responses to congenital viral infection

To explore how gestational timing shapes maternal and fetal antiviral immunity, we investigated maternal and fetal responses to congenital viral infection during early and mid-gestation using our previously established mouse model of congenital OROV infection^15^. In brief, we generated timed pregnant mice (dams) by mating C57BL6 (wild type; WT) 9-12 week-old female mice with WT males overnight and checking for the presence of a copulation plug the following morning, demarking embryonic day 0.5 (E0.5). Thereafter, 24 hours prior to the start of infection (early gestation: E6.5; mid-gestation: E11.5) pregnant mice were injected with 0.5 mg of IFNAR1 receptor (αIFNAR1)-blocking (or isotype control) antibody. Subsequently, pregnant dams were infected with OROV (prototype strain TR9760) (Fig. 1A). Broadly, our experimental design allows for direct comparison of fetal tissues from early and mid-gestation-infected dams matched to either gestation day or day post-infection (dpi) (Fig. 1A).

**Figure 1.**
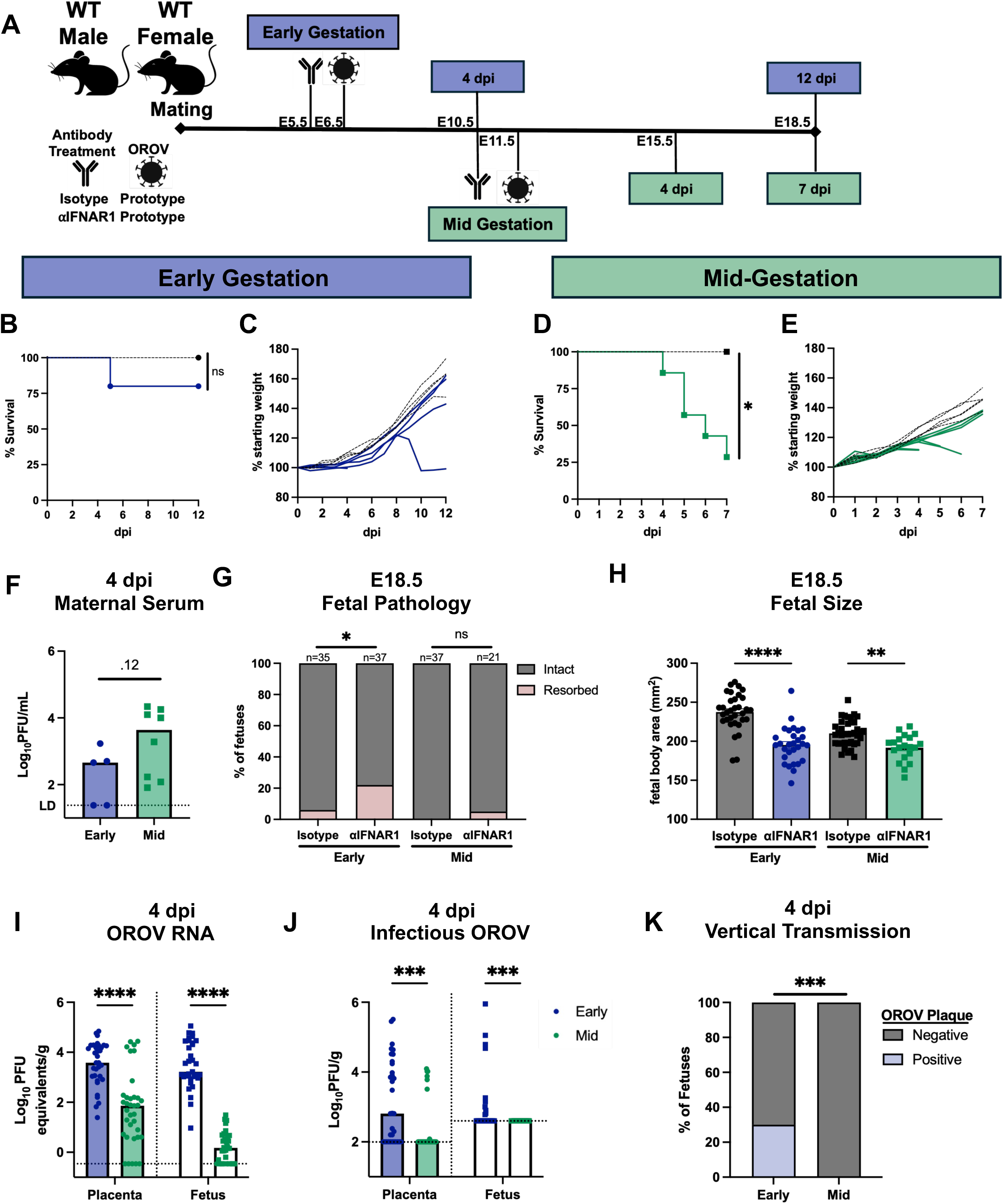
Discordant maternal and fetal responses occur across gestational stage during congenital viral infection. **A.** Experimental schematic depicting the timeline of both early gestation and mid-gestation congenital infection with OROV with corresponding embryonic day/day post-infection (dpi) indicated. **B-H.** C57BL/6 WT dams were pretreated with isotype control or αIFNAR1 and infected with OROV at early or mid-gestation through E18.5 in which we evaluated fetal gross pathology. **B.** Survival curve and percentage of starting weight **(C)** of mice in early gestation pretreated with either isotype control (dotted black line, n=4) or αIFNAR1 (blue line; n=5) and infected with OROV through 12 dpi. **D.** Survival curve and percentage of starting weight **(E)** of mice in mid-gestation pretreated with either isotype control (dotted black line, n=5) or αIFNAR1 (green line, n=8) and infected with OROV through 7 dpi. Statistical differences determined by Mantel-Cox test at either early or mid-gestation, *=p < 0.05, ns= not significant. **F.** Infectious OROV viral loads measured by plaque assay at 4 dpi in serum from the αIFNAR1-pretreated dams infected with OROV during early or mid-gestation. Statistical analysis performed via a Mann-Whitney *U* Test, p= 0.12. **G.** The rate of fetal resorptions on E18.5 across litters from isotype-control or αIFNAR1-treated pregnant mice infected with OROV at either early or mid-gestation. Statistical differences measured by Fisher’s exact test, *=p < 0.05, ns= not significant. **H.** Fetal size, measured by crown-rump length multiplied by occipital diameter, at E18.5 from isotype-control or αIFNAR1-treated pregnant mice infected with OROV at either early or mid-gestation. Statistical differences measured by Student’s T test, ****=p < 0.0001, **= p<0.01. **I-K.** Early (n=8) or mid-gestation (n=7) dams were pretreated with αIFNAR1, infected with OROV, and placental and fetal tissues were harvested at 4 dpi. **I.** OROV RNA, measured by RT-qPCR, and infectious OROV **(J)**, measured by plaque assay, from isolated placental or fetal tissue from early gestation-or mid-gestation-infected dams at 4 dpi. Statistical analysis performed via a Mann-Whitney *U* Test, ****=p<0.0001, ***=p<0.001. **K.** The rate of productive vertical transmission, determined by the presence or absence of plaquable OROV in fetuses from early or mid-gestation-infected dams at 4 dpi. Statistical differences measured by Fisher’s exact test, ***=p < 0.001. Dotted lines at the bottom of graphs denote the limit of detection in each assay.

As expected from our previous work^15^, dams pretreated with an isotype control antibody and infected with OROV during early gestation survive infection (Fig. 1B) with no signs of disease through 12 dpi (Fig. 1C). Further, dams pretreated with αIFNAR1 and infected with OROV during early gestation predominantly survive infection (Fig. 1B) and exhibit modest signs of disease (Fig. 1C). Interestingly, one dam experienced complete litter loss, as evidenced by weight loss between 8-10 dpi, despite exhibiting no other signs of disease (Fig. 1C), highlighting the propensity of OROV infection to adversely impact pregnancy outcomes even in the absence of overt maternal disease. Strikingly, when we performed similar experiments at mid-gestation, maternal disease was substantially more severe upon congenital OROV infection. We found most αIFNAR1-dams infected with OROV during mid-gestation succumb to infection by 7 dpi (Fig. 1D; 5 of 8 dams succumb) with considerable signs of disease (Fig. 1E). Notably, one dam exhibiting severe disease required euthanasia following premature delivery at E16.5; however, examination of the fetuses revealed no overt pathology (Fig. 1E). Differences in gestational-stage-dependent maternal susceptibility to severe OROV disease with trending, but statistically insignificant differences, in serum viral loads at 4 dpi (E10.5 or E15.5 for early or mid-gestation dams, respectively) (Fig. 1F). Overall, these data indicate gestational stage contributes to substantial differences in maternal morbidity and mortality upon OROV infection.

Interestingly, evaluation of gross fetal pathology at E18.5 revealed outcomes that were discordant with maternal disease severity across gestational stages (Supplementary Figure 1; representative litter photos). Although dams infected during early gestation were largely resistant to severe OROV disease, the rate of fetal resorption (dying tissue comprised of fetal and placental tissue) were significantly increased compared with those from isotype-treated controls (Fig. 1G). Conversely, fetal resorption rates among mid-gestation-infected dams were comparable to those of isotype-treated controls, despite severe maternal morbidity and mortality (Fig. 1G), suggesting that fetal outcomes were inversely associated with maternal disease severity. Further, this pattern was further reflected within the subset of intact fetuses in that intact fetuses from dams infected during early gestation were significantly smaller than the intact fetuses from matched isotype control-treated dams (Fig. 1H). In contrast, fetal size was only modestly, although significantly, reduced following infection during mid gestation (Fig. 1H), further supporting the observation that fetal pathology becomes increasingly restricted as gestation progresses despite more severe maternal disease.

We next asked whether differences in fetal pathology were a result of direct control of viral infection or a consequence of fetal immunopathology. At 4 dpi, there was a ∼1000x reduction in OROV RNA from early to mid-gestation fetuses (Fig. 1I), corresponding to a reduction in infectious OROV in fetuses from early compared to mid-gestation-infected dams (Fig. 1J). To this end, we found a complete block of productive OROV vertical transmission in fetuses from mid-gestation-infected dams, whereas 30% of fetuses from early gestation-infected dams were productively infected by OROV at 4 dpi (Fig. 1K). These data suggest that improved fetal pathology as gestation progresses may depend on the ability of the fetus to directly control vertical transmission.

### Fetal tissues clear viral infection independent of gestational stage

Given the discordance between maternal disease and fetal outcomes across gestation, we next asked whether fetal control of OROV throughout infection differs with gestational stage. To do this, we analyzed OROV RNA and infectious virus in distinct fetal tissues (placenta, fetus, or resorptions) from early and mid-gestation-infected dams over the course of infection (12 or 7 dpi, respectively) until one day prior to birth (E18.5). Upon OROV infection during early gestation, placentas and intact fetuses cleared OROV RNA (Fig. 2A) and infectious virus (Fig. 2B) as infection progressed from 4 to 7 to 12 dpi. Similarly, placentas and fetuses from mid gestation-infected dams also cleared OROV RNA and infectious virus, highlighting the shared ability of fetal tissues to clear OROV infection across gestational stage (Fig. 2C-2D). Interestingly, upon infection during early gestation, we noted that the fetal units that failed to clear OROV RNA by E18.5 corresponded to resorbed fetuses, evidenced by resorptions having significantly more viral RNA than their intact fetal counterparts (Fig. 2E). Although our interpretations are confounded by the infrequent instance of resorptions upon mid gestation infection, we detected more OROV RNA in the sole resorption from a mid-gestation-infected dam than its intact fetal counterparts (Fig. 2E). Altogether, these data indicate that fetal control and clearance of OROV is independent of the gestational stage and correlates with improved fetal pathology.

**Figure 2.**
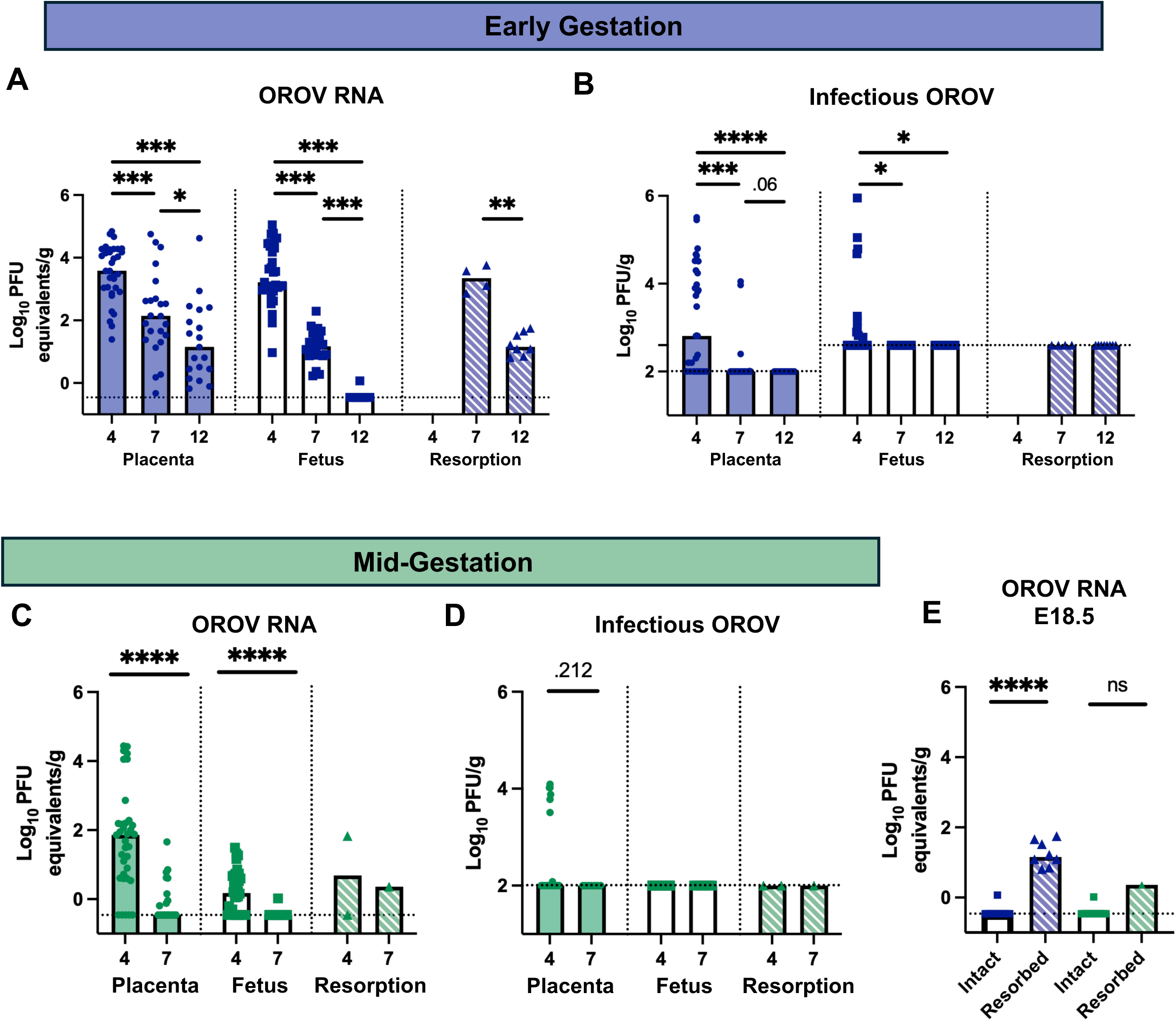
Fetal tissues independent of gestational stage clear OROV infection. A-E. Dams were pretreated with αIFNAR1 and infected with OROV either during early gestation or mid-gestation, and fetal tissues were collected at indicated timepoints. **A.** OROV RNA or infectious OROV (**B**) was quantified at 4 dpi (8 dams), 7 dpi (3 dams; reanalysis of data published in PMID: 40964022), and 12 dpi (4 dams) from the placenta, intact fetus, or resorbed fetal unit (placental and fetal tissue) from early gestation-infected dams. **C.** OROV RNA or infectious OROV (**D**) was quantified at 4 dpi (7 dams) and 7 dpi (3 dams) from placenta, intact fetus, or resorbed fetal unit from mid-gestation-infected dams. **E.** OROV RNA from intact fetuses compared to resorbed fetal tissues at E18.5 from both early and mid-gestation infected dams. Statistical analysis performed via a Mann-Whitney *U* Test with multiple comparison corrections, ****=p<0.0001, ***=p<0.001 *=p<0.05, ns=p>0.05. Dotted lines at the bottom of graphs denote the limit of detection in each assay.

### Differential susceptibility of distinct regions across the maternal-fetal interface reveals potential routes of OROV vertical transmission

We next sought to identify the regions of the maternal-fetal interface that are susceptible to OROV infection and may facilitate vertical transmission. Because we did not observe productive vertical transmission to fetuses following mid-gestation infection by 4 dpi, we focused our analysis on E10.5 (4 dpi) implantation sites from early gestation-infected (Fig. 1A) when vertical transmission was readily detected (Fig. 1J). We sectioned and analyzed whole E10.5 implantation sites to maintain the natural architecture of the entire fetal unit and delineated five anatomically distinct regions across the maternal-fetal interface. Based on established tissue-specific markers and spatial organization, these included, from left to right in Figure 3A: contralateral uterus (relative to the fetal placenta), fetus, fetal placenta, decidua, and ipsilateral uterus (relative to the placenta). By evaluating 20 fetuses across 7 independent dams, we observed reproductible and robust OROV antigen across multiple regions of the maternal-fetal interface, with similar patterns between distinct implantation sites and across independent litters, whereas implantation sites from mock-infected dams lacked detectable OROV signal in all regions (Fig 3B). We found abundant and increased OROV infection, represented by increased cumulative total area of OROV staining, in the ipsilateral uterus compared to fetal and maternal regions of the placenta (Fig. 3C). Strikingly, however, we found the contralateral uterus to be ∼5x more infected with OROV by total area compared to the ipsilateral uterus (Fig. 3C). Comparatively, the fetal placenta and the fetus had far less total OROV staining by area despite being adjacent to highly infected maternal uterine regions, suggesting that fetal immune mechanisms are inhibiting OROV transmission from maternal to fetal tissues at these regional interfaces (Fig. 3C). Additionally, we noted no differences in OROV staining between the fetus, fetal placenta, and decidua (Fig.3C), suggesting that placenta-specific immune responses may be occurring locally to protect both maternal and fetal placental tissue from OROV infection. Analysis of individual implantation sites across litters revealed considerable heterogeneity in OROV infection (Fig. 3D), consistent with our quantification of OROV RNA and infectious virus in fetal tissues. (Fig. 1I-J)).

**Figure 3.**
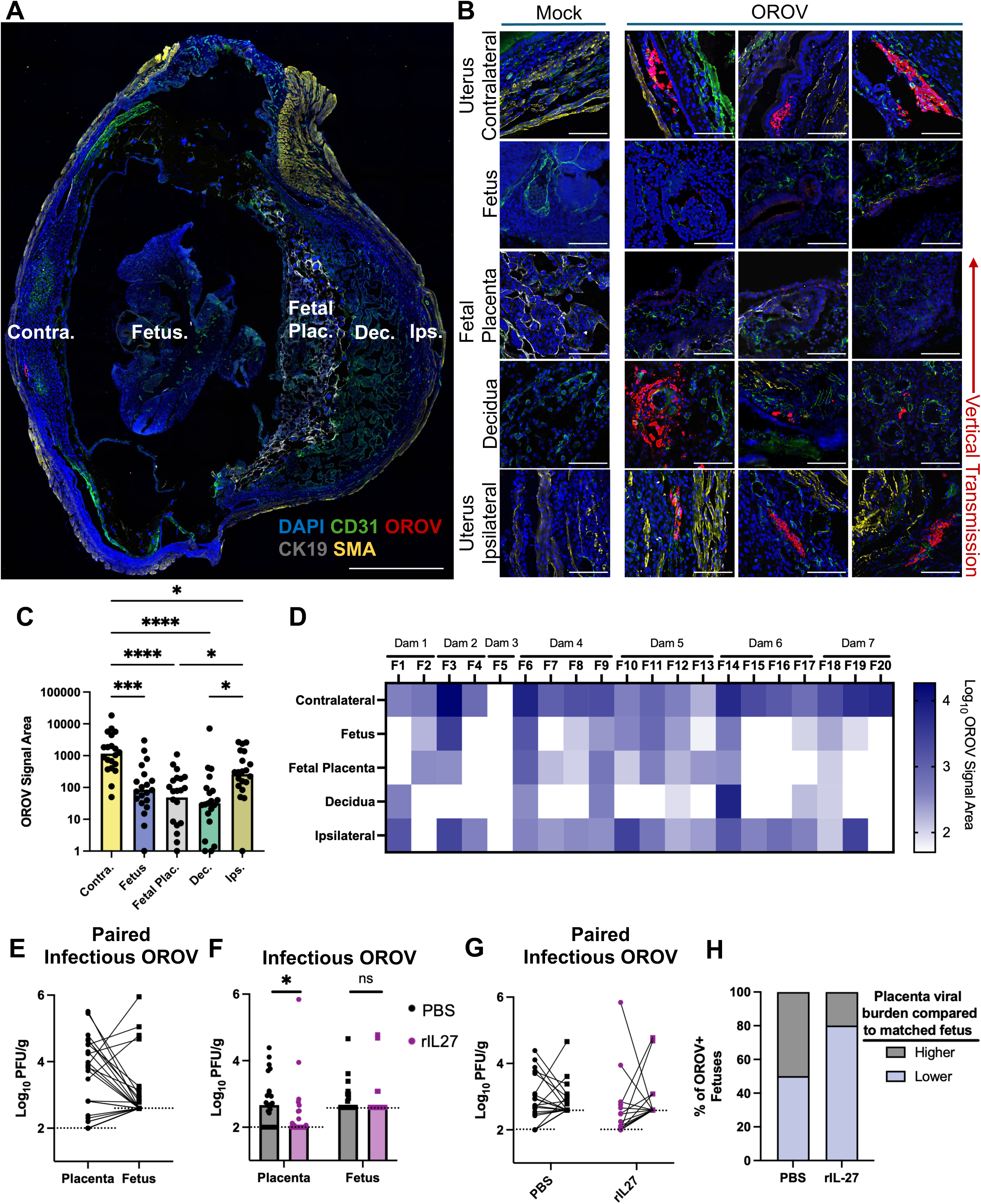
Distinct regions of the maternal-fetal interface are differentially susceptible to viral infection and may contribute to vertical transmission. A-D. Dams (n=7) were pretreated with αIFNAR1 and infected with OROV during early gestation, and implantations sites were collected at 4 dpi. **A.** Representative immunofluorescent image of whole murine implantation site at gestational day E10.5. Five distinct anatomical regions of the murine implantation site (Contra., Fetus., Fetal Plac., Dec., Ips.) were determined via immunofluorescent staining of endothelial cells (CD31, green), viral antigen (OROV, red), trophoblasts (CK19, gray), smooth muscle actin (SMA, yellow) and nuclei (DAPI, blue). Contra: contralateral uterus; Ips: ipsilateral uterus; Dec: decidua; Fetal Plac: Fetal Placenta. 20X magnification, stitched. Scale bar, 1000μm. **B.** Representative immunofluorescent images of mock-and OROV-infected implantation sites at gestational day E10.5, by anatomical region. For each region, selected images are representative of 3 different implantation sites from 3 independently infected dams. 20x magnification, zoomed. Scale bar 100 μm. **C.** Quantification of OROV signal area (microns^2^) in distinct anatomical regions of murine embryo (Contra., Fetus., Plac., Dec., Ips.), as determined by ImageJ software. Graph represents 20 implantation sites from 7 independent litters, with at least 9 images acquired per region per implantation site. Statistical analysis performed via one-way ANOVA adjusted for multiple comparisons, ****= p<0.0001, ***= p<0.001, *=p<0.05. **D.** Heatmap displaying the Log_10_ value of OROV signal area of OROV staining across each region within each implantation site. Dark blue represents the most abundantly stained regions while white represents regions without OROV staining. **E.** Paired analysis between matched placental and fetal infectious OROV viral loads from early gestation-infected dams at 4 dpi in Figure 2B. **F-H.** WT dams infected during early gestation treated with either PBS (n=4) or recombinant IL-27 (n=4) at 0 and 2 dpi and placental and fetal tissue collected for viral load quantification at 4 dpi. Unpaired (**F**) and paired (**G**) scatter plots of infectious OROV measured by plaque assay. **H.** Categorical representation of paired placental and fetal viral loads depicting the incidence in which placental viral loads were higher or lower than viral loads in matched fetal samples from PBS-or rIL-27-reated dams. Statistical analysis in **F** performed via a Mann-Whitney *U* Test, *=p<0.05. Dotted lines at the bottom of graphs denote the limit of detection in each assay.

Overall, OROV antigen signal area decreases between ipsilateral uterus, placental and fetal tissue, representing the expected bottleneck of transplacental transmission (Fig. 3C-3D). However, we found some cases in which the fetus had substantial OROV infection despite reduced or absent infection in the maternal decidua or fetal placenta (Fig. 3D; ie. F3, F19). Interestingly, cases of increased fetal infection compared to its respective placenta were accompanied by abundant OROV staining in the maternal contralateral uterus (Fig. 3D; F3, F19). These data suggest that, in some instances, maternal-to-fetal OROV transmission may occur through routes that bypass extensive placental infection and implicate the contralateral uterus as a region that can potentially facilitate vertical transmission. Upon a paired inspection of our infectious OROV tissue data (Fig. 2B), we predominantly observed greater infection in the placenta compared to the matched fetal counterparts (Fig. 3E). Interestingly, however, we sporadically observed greater OROV infection in the fetus than in the corresponding placenta across paired samples (Fig. 3E) correlating with individual staining patterns observed via immunofluorescence (Fig. 3D). To further investigate whether fetal infection can occur independently of placental viral burden, we treated dams infected during early gestation with PBS or recombinant Fc-conjugated IL-27 (rIL-27) at 0 and 2 dpi. We have previously identified endogenous IL-27 as a mediator of placental, but not fetal, antiviral immunity against ZIKV and therefore used this approach to selectively enhance antiviral control within the placenta^7^. As expected, rIL-27 treatment significantly reduced placental OROV viral loads at 4 dpi, while fetal viral loads were unchanged compared to PBS-treated dams (Fig. 3F). Further, paired analysis identified fetuses with substantial OROV infection despite low or undetectable OROV infection in matched placentas (Fig. 3G). Additionally, upon IL-27 treatment we found a trending increase in the number of productively infected fetuses with lower viral burdens in their respective placentas compared to PBS-treatment (Fig. 3H), suggesting that by exogenously boosting placental antiviral immunity, we could reveal alternative routes of vertical transmission to the fetus independent of the placenta.

Altogether, these data reveal a mosaic pattern of viral replication across the maternal-fetal interface. OROV infection generally decreased stepwise from maternal to fetal tissues, consistent with increasing restriction of infection as the virus approaches the fetus and highlighting that fetal-intrinsic antiviral responses may limit vertical transmission. Moreover, the presence of substantial fetal infection despite minimal or undetectable infection in matched placentas raises the possibility of an alternative route of vertical transmission, potentially through contact with the contralateral region of the maternal uterus, that does not require extensive placental infection.

### Inter-fetal immune responses restrict vertical transmission and pathology in a type I IFN-dependent manner

Finally, we sought to determine which fetal antiviral immune responses contribute to restricting vertical transmission, replication, and pathology. Given the dependence of maternal infection upon transient IFNAR1 blockade in our murine infection model^15^, we asked whether fetal type I IFN signaling influenced OROV vertical transmission, clearance, and associated pathology. We employed a mating strategy in which heterozygous IFNAR1^+/-^ females were mated to either IFNAR1^-/-^ males (Fig. 4A, Cross A) or to WT males as a control (Fig. 4B, cross B) to generate mixed genotype litters. Thereafter, we pretreated with αIFNAR1 and infected heterozygous dams during early gestation and evaluated placental and fetal viral loads at 4 dpi. Surprisingly, we found no differences in placental and fetal infectious viral loads at 4 dpi between IFNAR1^-/-^ and IFNAR1^+/-^ fetuses from cross A nor in IFNAR1^+/+^ and IFNAR1^+/-^ fetuses from cross B (Fig. 4A-B), indicating that fetal IFNAR1 signaling does not directly control OROV vertical transmission or local replication within the fetus.

**Figure 4.**
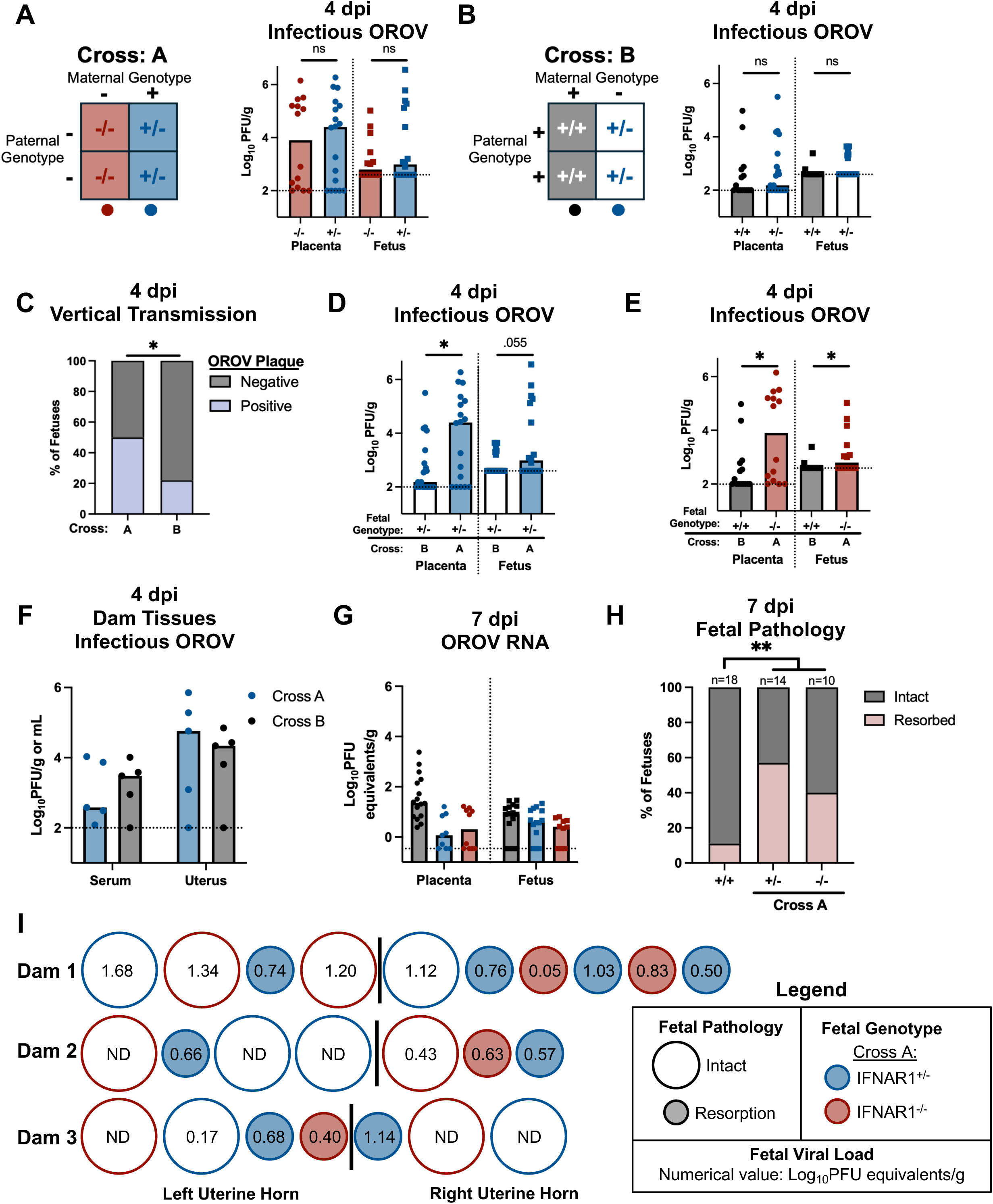
Inter-fetal antiviral immunity restricts vertical transmission and pathology in a type I IFN-dependent manner. A-F. IFNAR1^+/-^ dams crossed with IFNAR1^-/-^ or WT sires were pretreated with αIFNAR1 and infected at early gestation with OROV and maternal, placental, and fetal tissues were collected at 4 dpi and quantified by plaque assay. Data represent tissue from 5 dams per cross. **A.** Cross A Punnett square outlining maternal genotype, paternal genotype, and potential ratios of resultant fetal genotypes. Infectious OROV viral loads between IFNAR1^-/-^ and IFNAR1^+/-^ placental and fetal tissue from Cross A are depicted. **B.** Cross B Punnett square outlining maternal genotype, paternal genotype, and potential ratios of resultant fetal genotypes. Infectious OROV viral loads between IFNAR1^+/-^ and IFNAR1^+/+^ placental and fetal tissue from Cross B are depicted. Statistical analysis performed via a Mann-Whitney *U* Test, ns=p>0.05. **C.** Aggregate rates of vertical transmission at 4 dpi, determined by detecting plaquable OROV in the fetus, between all fetuses from Cross A compared to Cross B. Statistical differences measured by Fisher’s exact test, *=p < 0.05. **D.** Infectious OROV viral loads in IFNAR1^+/-^ placental and fetal tissue from Cross A compared to Cross B. **E.** Infectious OROV viral loads in IFNAR1^+/+^ placental and fetal tissue from Cross B compared to IFNAR1^-/-^ placental and fetal tissue from Cross A. Statistical analysis performed via a Mann-Whitney *U* Test, *=p<0.05 or p=0.055. **F.** Infectious OROV viral loads in the serum and uterus from Cross A and Cross B IFNAR1^+/-^ dams at 4 dpi. Statistical analysis performed via a Mann-Whitney *U* Test, values were ns=p>0.05. **G-H.** WT dams mated to WT sires (n=3) or IFNAR1^+/-^ dams mated to IFNAR1^-/-^ sires (Cross A, n=3) were infected with OROV during early gestation and evaluated through 7 dpi. **G.** OROV RNA was quantified in placental and fetal tissue by qPCR and (**H**) gross fetal pathology, determined by resorption rate, was evaluated at 7 dpi. Statistical analysis for OROV RNA performed via a Mann-Whitney *U* Test correcting for multiple comparisons, values were ns=p>0.05. Statistical differences measured by Fisher’s exact test between fetal pathology in Cross A compared to WT x WT fetuses, **=p < 0.01. **I.** Representation of individual fetal information in fetuses derived from each Cross A dam at 7 dpi depicting fetal pathology (shape), fetal genotype (color), or fetal viral load (number; ND= not detectable) as well as spatial organization of fetuses across the litter. The black vertical line denotes the left from the right uterine horn. Dotted lines at the bottom of graphs denote the limit of detection in each assay.

However, when we compared the incidence of OROV vertical transmission at 4 dpi between the fetuses derived from Cross A (IFNAR1^-/-^ and IFNAR1^+/-^ mixed litters) compared to the fetuses derived from Cross B (IFNAR1^+/+^ and IFNAR1^+/-^ mixed litters), we observed striking differences (Fig. 4C). There was a significantly greater incidence of vertical transmission among fetuses from Cross A, with 50% of fetuses OROV+ by plaque assay, compared with only 22% from Cross B. These findings suggest that while there is not a direct role for IFNAR1 signaling in individual fetuses at 4 dpi, IFNAR signaling may contribute to inter-fetal immunity to control OROV vertical transmission. To further evaluate this possibility, we compared genetically identical IFNAR1^+/-^ placental and fetal tissues derived from either OROV-infected heterozygous IFNAR1^+/-^ dams from Cross A or Cross B. We observed substantially increased infectious OROV in both IFNAR1^+/-^ placental and fetal tissue from Cross A compared to IFNAR1^+/-^ placental and fetal tissue from Cross B (Fig. 4D). Because these IFNAR1^+/-^ fetuses from Cross A and Cross B differ only by the IFNAR1 genotype of the fetuses with which they share an intrauterine environment, these findings suggest that the immune capacity of neighboring fetuses influences susceptibility to OROV infection and vertical transmission, supporting a role for inter-fetal antiviral immunity.

We next compared infectious viral burdens in homozygous WT (IFNAR1^+/+^) placental and fetal tissues from Cross B with homozygous IFNAR1^-/-^ tissues from Cross A. We observed substantially and significantly increased infectious OROV in IFNAR1^-/-^ tissue from Cross A compared to IFNAR1^+/+^ placental and fetal tissue from Cross B (Fig. 4E), further supporting a role for fetal type I IFN signaling in inter-fetal immune responses to OROV by 4 dpi. To determine whether differences in maternal infection between the two crosses could contribute to these fetal outcomes, we measured infectious OROV burden in serum and uterine tissues of Cross A and Cross B dams at 4 dpi. Viral burdens were comparable between the two groups (Fig. 4F), suggesting that the differences in fetal infection were not attributable to altered maternal viral burden. Altogether, these findings support an IFNAR1-dependent mechanism of inter-fetal antiviral immunity that restricts OROV vertical transmission.

We next asked whether type I IFN-dependent inter-fetal immunity also contributed to fetal OROV clearance and pathology. Given mouse availability and the absence of differences in viral burdens associated with heterozygous IFNAR1^+/-^ dams or fetuses (Fig. 4B), we compared OROV RNA burden and fetal pathology at 7 dpi between Cross A and WT x WT matings. We found no differences in OROV RNA burden among WT, IFNAR1^-/-^, and IFNAR1^+/-^ placental or fetal tissues, suggesting that fetal clearance of OROV RNA occurs independently of type I IFN signaling (Fig. 4G).

Despite comparable viral burdens, we observed striking differences in fetal pathology between WT x WT compared to Cross A fetuses at 7 dpi (images of Cross A pathology in Supplementary Figure 2A). Whereas only 11% of WT x WT fetuses are resorbed, 57% of IFNAR1^+/-^ fetuses and 40% of IFNAR1^-/-^ fetuses were resorbed by 7 dpi (Fig. 4H). Among intact fetuses, however fetal size did not differ across all genotypes or crosses (Supplementary Fig. 2B). Notably, we observed spatial clustering of resorptions across Cross A fetuses (Fig. 4I; Supplementary Fig. 2A). These clusters contained both IFNAR1^-/-^ and IFNAR1^+/-^ fetuses and were not associated with differences in viral burden (Fig. 4I), suggesting that neither fetal genotype nor viral load alone explained the spatial pattern of pathology. Altogether these findings support a model where type I IFN signaling is dispensable for eventual fetal clearance of viral RNA but coordinates inter-fetal antiviral immunity that protects against severe fetal pathology.

Overall, our data support a model of fetal antiviral immunity in which inter-fetal communication contributes to resistance to vertical transmission and severe fetal pathology. Although, we found no evidence for a direct role of fetal IFNAR signaling in determining vertical transmission by 4 dpi or fetal pathology at 7 dpi, our findings reveal a broader requirement for type I IFN signaling in coordinating inter-fetal immune responses. Moreover, spatial clustering of fetal resorptions, independent of individual fetal genotype or viral burden, suggests that protection may be spatially coordinated within the intrauterine environment. Together, these findings uncover an IFNAR-dependent, inter-fetal antiviral immune response that extends beyond the immune capacity of an individual fetus to protect fetal health.

## DISCUSSION

In this study, we sought to define how maternal and fetal immune responses changed across gestation. We found that maternal immune responses during early gestation were highly effective at controlling OROV infection and disease, particularly compared with responses during mid-gestation (Fig. 1). This is distinct from literature using ZIKV, in which gestational stage-dependent differences in maternal viral disease have not been observed^2^. Despite being separated by only 4 days of gestation, these distinct maternal outcomes underscore the dynamic nature of immunity across pregnancy and reveal a substantial shift in the maternal immune environment with important consequences for maternal health. Mid-gestation is characterized by a more tolerogenic immune state that supports the development of the antigenically distinct fetus^1^, which may come at the cost of dampened systemic antiviral immunity against pathogens such as OROV. Our findings in Figure 1 highlight the need to further define how maternal antiviral immunity changes across gestation and how these changes shape susceptibility to infection.

Maternal and fetal outcomes are strikingly discordant during OROV infection. While maternal outcomes are substantially better following infection during early gestation, fetal pathology is markedly increased compared with fetuses from dams infected during mid-gestation, despite considerable maternal morbidity and mortality (Fig. 1). Consistent with these divergent fetal outcomes, mid-gestation placental and fetal tissue harbor 100-1000-fold lower levels of OROV RNA than during early gestation infection (Fig. 1I). Moreover, we observed a complete block in productive vertical transmission to the fetus during mid-gestation, whereas productive vertical transmission occurred in approximately 30% of fetuses following early gestation infection (FIg.1J-1K). These findings suggest that antiviral mechanisms emerge over the course of fetal development that more effectively restrict vertical transmission and protect fetal health, even in the setting of severe maternal disease. Altogether, our data underscores the dynamic nature of fetal antiviral immunity across gestation and highlights the fetus as an active contributor to protection from congenital viral infection.

We report that the placenta and fetus are not only capable of restricting OROV infection but can also clear infection over time. To our knowledge, other congenital viral infection models generally exhibit persistent or increasing viral burdens in fetal tissues as infection and gestation progress. In contrast, we observed robust clearance of both infectious OROV and viral RNA from placental and fetal tissue, regardless of gestational stage (Fig. 2). Many studies of congenital viral infection, including congenital OROV infection, rely predominantly on viral RNA quantification by qPCR to measure viral burden in fetal tissue^15–18^. By measuring both infectious viral by plaque assay and viral RNA by qPCR, our study reveals distinct infection dynamics captured by each approach and demonstrates that detection of viral RNA does not necessarily reflect ongoing productive infection. Notably, by E18.5, the only fetal units in which OROV RNA remained detectable had undergone resorption, linking failure to clear viral RNA with severe fetal pathology (Fig. 2E). Altogether these data suggest fetal intrinsic antiviral mechanisms acting after the initial vertical transmission event shape subsequent infection outcomes, promoting either viral clearance and fetal protection or persistent infection and severe fetal pathology.

Our study provides the first extensive spatial characterization of OROV infection by immunofluorescence across the maternal-fetal interface. In aggregate, we observed substantially greater OROV staining in the ipsilateral region of the maternal uterus compared to the decidua, fetal placenta or fetus, revealing a stepwise decrease in infection from maternal to fetal tissues (Fig. 3C). By evaluating regional OROV infection dynamics across 20 implantation sites from 7 independent dams during early gestation, we further uncovered considerable spatial heterogeneity in infection throughout maternal and fetal reproductive tissues. Notably, several of the most highly infected fetuses were positioned adjacent to heavily infected regions of the contralateral uterus, despite comparatively limited infection of their corresponding placentas (Fig. 3D). These observations raise the possibility that OROV can transmit directly from the contralateral uterus to fetal tissues, serving as a placental-independent route of vertical transmission. Similar routes have been proposed in mouse and nonhuman primate models of congenital ZIKV infection, in which close contact between maternal uterine and fetal tissues may facilitate viral transmission^19–21^. Such a mechanism could provide a means for a virus to bypass potent antiviral defenses within the placenta and directly access the fetus. Although additional studies are required to experimentally establish this route of transmission, our spatial analyses offers compelling evidence implicating the contralateral uterus as a potential contributor to OROV vertical transmission (Fig. 3).

Finally, we uncovered a unique and previously unreported mechanism of fetal antiviral immunity. Using heterozygous-by-homozygous mating schemes, we found no evidence that fetal-intrinsic IFNAR signaling directly controls OROV vertical transmission or fetal pathology (Fig. 4). This finding is consistent with earlier models of congenital OROV infection in which fetal IFNAR signaling did not substantially alter infection outcomes^17^. Surprisingly, however, comparison across our two mating schemes revealed that the broader fetal IFNAR environment substantially influenced both vertical transmission and fetal pathology (Fig. 4C-E, 4H). Specifically, genetically identical IFNAR1+/-fetuses exhibited markedly different outcomes depending on whether they developed alongside IFNAR1+/+ or IFNAR1-/-littermates, suggesting that type I IFN signaling contributes to inter-fetal immune crosstalk. One potential mechanism is that robust OROV infection in the absence of IFNAR signaling initiates a local inflammatory cascade that alters the immune environment of neighboring fetuses, thereby increasing susceptibility to vertical transmission and severe fetal pathology. Alternatively, differences between the two crosses could arise from paternal genotype through mechanisms independent of the resultant fetal genotypes. We are particularly intrigued by the possibility given our recent findings demonstrating nongenetic paternal contributions to offspring immune outcomes^22^. However, more experimental work is needed to test these hypotheses. Nevertheless, our findings reveal that fetal antiviral immunity may extend beyond the individual fetus, with protection spatially coordinated across the intrauterine environment through IFNAR-dependent inter-fetal immune responses that influence vertical transmission and congenital pathology.

Altogether, our group employs an emerging congenital pathogen to uncover fetal-intrinsic antiviral mechanisms that emerge over the course of gestation. We demonstrate that maternal and fetal outcomes during OROV infection can be strikingly uncoupled across gestational stages and link gestation-dependent differences in fetal pathology to susceptibility to vertical transmission. Remarkably, we find that fetal tissues can actively clear viral infection over time, regardless of gestational stage, and identify regional differences in antiviral control across the maternal-fetal interface that may shape routes of vertical transmission. Finally, we reveal a previously unrecognized dimension of fetal antiviral immunity in which inter-fetal immune crosstalk contributes to the control of OROV vertical transmission and associated fetal pathology. Collectively, these findings establish the fetus as an active participant in antiviral defense and reveal that fetal immunity is dynamic, spatially organized, and coordinated across the intrauterine environment. Our work provides a framework for understanding how fetal antiviral immunity develops across gestation and may ultimately inform strategies to protect fetal health during congenital viral infection.

## MATERIALS AND METHODS

### Cells and viruses

OROV strain TR9760 (prototype) was purchased from ATCC (VR-266). OROV stocks were grown from the initial isolate on Vero cells as a P1 stock. P1 stocks of each virus were then grown again on Vero cells and concentrated by ultrafiltration (Centricon Plus-70; MWCO: 30,000) to generate a P2 stock for experimental use. African green monkey kidney (Vero) (ATCC, Cat#CCL-81) cells were grown and maintained in Dulbecco’s modification of eagle’s medium (Corning, Cat#10–013-CV) supplemented with 5% fetal bovine serum (Sigma-Aldrich, Cat#F2442) and 1% penicillin–streptomycin (ThermoFisher Scientific, Cat#15140122). Cell lines were authenticated by morphology and were routinely tested for mycoplasma contamination.

### Mice

C57BL/6 (wild-type; WT) and IFNAR1^-/-^ mice were purchased from The Jackson Laboratory (strain #000664 and strain #028288, respectively) and maintained at the University of Pennsylvania under specific pathogen-free conditions. IFNAR1^+/-^ mice were generated and maintained in house by breeding a IFNAR1^-/-^ male with a WT female. All timed matings were performed by mating >8-week old female mice to male mice overnight. Subsequently male and female mice were separated the following morning after checking for the presence or absence of a copulation plug, with plug observation date designated E0.5. Female mice with a copulation plug were used for congenital OROV infection experiments. All experiments were performed following IACUC guidelines.

### Congenital OROV infection

9–12-week old pregnant female mice were treated intraperitoneally with 0.5mg/mouse of either an αIFNAR1-blocking antibody (clone MAR1–5A3; BioXCell Cat#BE0241) or an IgG1 isotype-control antibody (BioXCell Cat#BE0083) on E5.5 for early gestation infections and E10.5 for mid-gestation infections. Treated pregnant mice were then infected subcutaneously via footpad injection with 10^4^ PFU of OROV on E6.5 (early gestation) or E11.5 (mid gestation). Mice were monitored daily for weight changes, overt disease signs, and sacrificed at indicated timepoints. Tissues were isolated, placenta and fetuses were photographed on measurement paper using an iPhone camera, assessed for gross pathology. Fetuses were defined as “intact” if they had an intact placenta and no clear signs of pathology. Fetuses were defined as “resorption” if they had clear loss of tissue integrity (in the presence or absence of a placenta) or lacked a distinguishable fetal and placental unit altogether. All tissues were collected into tubes containing 1mL of DMEM with 2%FBS, 1% penicillin/streptomycin, 1% HEPES + silica beads, homogenized, and analyzed for subsequently for viral loads. Of note, bars representing 7 dpi placental, fetal, and resorption viral loads by OROV RNA (Fig. 2A) and infectious OROV (Fig. 2B) are a reanalysis of data previously published in our initial manuscript reporting the congenital OROV infection model^15^.

### Fetal size quantification

Fetal size was quantified using photos of respective litters and converting pixels to millimeters (mm) via reference graph paper within each photo. For intact fetuses, fetal crown-rump length, the direct length from the crown of the fetal head to the start of the tail, was multiplied by occipital diameter, the direct length from the front (near the eye) to the back of the fetal head, to quantify overall fetal size. All analysis was performed using ImageJ software.

### Plaque assay

Vero cells were seeded at a density of 7.5×10^5^ cells/well in a 6-well plate overnight. After 24-hours, sample homogenates (or serum) were serially diluted in DMEM with 2% FBS, 1% penicillin/streptomycin, 1% HEPES. Seeded Vero cells were treated with 250μL of diluted supernatants and incubated for 1-hour, with rotations every 15 minutes. Following incubation, inoculum was aspirated and replaced with MEM (Sigma-Aldrich, Cat#11430030) supplemented with 5% FBS, 1% GlutaMAX, 1% Non-essential amino acids, and 0.65% agarose (Lonza, Cat#50111). Plates were fixed at 4 dpi with 2 mL 10% NBF (ThermoFisher Scientific, Cat#22050105) and visualized using 0.1% crystal violet (ThermoFisher Scientific, Cat#C581– 25). Plaques were manually counted, and virus titer was calculated.

### RNA Extraction, cDNA synthesis, and qPCR

OROV RNA in infected tissues was quantified by adding 200μL of tissue homogenate or serum to 600μL of TRIzol and then frozen at −80. RNA was then subsequently extracted using Phasemaker (Invitrogen; #A33251) tubes according to manufacturer’s protocol. Total RNA was then quantified using the ThermoFisher Scientific Nanodrop One spectrophotometer and cDNA was generated as above using 500ng total RNA per reaction using an iScript cDNA synthesis kit (Bio-Rad, Cat#1708890) according to manufacturer’s instructions. OROV cDNA was quantified by quantitative PCR with Power SYBR Green Master Mix (ThermoFisher Scientific, Cat#4367659). Reactions were run via QuantStudio3 (50 °C: 2’; 95 °C: 10’; 40 × 95 °C: 15 s, 60°C: 1’) with the addition of a final melt curve (95 °C: 15 s; 60 °C: 1’; 95 °C: 1’). All samples were loaded in technical duplicates. OROV cDNA was specifically amplified using previously published primers targeting the S segment of OROV (F: 5′-GCGTCACCATCATTCCAAGTA-3′, R: 5′-CCCAGATGCGATCACCAATTA-3′)^15,16^. Ct values were fitted to a standard curve generated from a known concentration (PFU/mL) of stock OROV-P to determine the PFU equivalents/mL of a given sample. Samples were then normalized to respective starting mass (g) or volume (mL) and reported as PFU equivalents/g or mL.

### Fetal IFNAR1 genotyping

Genomic material from E10.5 fetal units is limited, and we observed considerable contamination of maternal heterogenous genetical material during dissection to confound accurate gel-based genotyping. To limit the amount of genomic material necessary for both viral load quantification by qPCR and genotyping, we designed a qPCR assay for genotyping using previously synthesized sample cDNA. Therefore, we designed 3 primer sets for quantifying the wild-type (containing exon 3) and mutant (lacking exon 3) *Ifnar1* transcripts by qPCR in 3 separate reactions: a positive control primer set nesting in exon 11, a wild-type transcript primer set nesting in exon 2 and exon 3, and a mutant transcript primer set nesting in exon 2 and exon 4.

IFNAR1^-/-^ fetuses were differentiated from IFNAR1^+/-^ fetuses by a delta CT between wild-type and control transcript amplification as >3.32 CT. IFNAR1^+/+^ fetuses were differentiated from IFNAR1^+/-^ fetuses by a delta CT between mutant and control transcript amplification as >3.32 CT. The primer sets are listed as follows: *Ifnar1* control (F: 5’-CTGAGGAGCACACGGAAAGA - 3’, R: 5’-AGGCGCGTGCTTTACTTCT-3’), *Ifnar1* wild-type (F: 5’-CGAACAAAAGACGAGGCGAA-3’, R: 5’-TCCTCTGCTCTGACACGAAAC-3’), *Ifnar1* mutant (F: 5’-GGCAGTGTGACCTTTTCAGC –3’, R: 5’-CGGGAGGAGAGATGTGGACT –3’).

### *In vivo* recombinant IL-27 treatments

Pregnant WT mice were treated with 25 µg (in 200µL of PBS) of recombinant-conjugated mouse IL-27 (or PBS vehicle control) intraperitoneally on 0 and 2 dpi. Recombinant Fc-conjugated mouse IL-27 was supplied by Dr. Christopher Hunter (University of Pennsylvania) and generated by Dr. Aaron Ring (Fred Hutchinson Cancer Center) according to published protocols^23^.

### Immunofluorescence

E10.5 mouse implantation sites were washed in PBS upon tissue collection and fixed in 4% paraformaldehyde (Cat#) in PBS overnight at 4°C. All tissues were then submerged in a 30% sucrose solution prior to cryopreservation in OCT tissue embedding medium (Sakura, 4583). 10μm cryosections of whole mouse implantation sites were then obtained using a Leica Cryostat. Thereafter, we then used 3 cryosections encompassing an early section, middle section, and late section for subsequent staining procedures. All cryosections were washed three times with 1x PBS prior to membrane disruption with PBS supplemented with 0.4% Triton X-100 (PBST; Sigma-Aldrich, T8787) and blocking with 2% normal goat serum (Sigma-Aldrich, G9023-5ML) in PBST for one hour at room temperature. Immunofluorescence was then performed with the following primary antibodies diluted in blocking solution: Armenian hamster anti-CD31 (1:10, Developmental Studies Hybridoma Bank, 2H8s), rat anti-Cytokeratin 19 (1:500, Abcam, EP1580Y, Cat#ab323561), anti-Smooth Muscle Actin Alexa Fluor Plus 750 (1:200, Invitrogen,1A4, Cat#757-9760-82) and rabbit anti-OROV N (1:2000, Custom Genescript gifted by Dr. Anita McElroy at the University of Pittsburgh). All slides were washed with three times with PBST, followed by incubation with secondary antibodies (Jackson ImmunoResearch) and 1x DAPI (ThermoFisher, AC20271010) diluted in PBST for 2 hours at room temperature. To reduce background signal intensity, whole embryo sections were subsequently incubated in TrueBlack Lipofuscin Autofluorescence Quencher (Biotium, 23007) for one minute at room temperature. All slides were then mounted with media (ThermoFisher, P36961) and a coverslip. Immunofluorescence images were acquired using a Nikon Eclipse Ti2 microscope and processed using NIS-Elements and ImageJ software. Total OROV signal area in murine implantation site images was quantified using the ImageJ particle count application and by totaling the signal area across early, middle, and late sections. Each value was then corrected by subtracting the cumulative total signal area from mock-infected implantation sites that were stained on the same day from the cumulative total signal area from OROV-infected implantation sites. For each implantation site, distinct anatomical regions of fetus, placenta, decidua, contralateral uterus and ipsilateral uterus were identified during imaging using previously established markers as well as spatial orientation. Briefly, the uterus was defined via SMA+/CD31+/CK19-cells, with the ipsilateral uterus located adjacent to the decidua and placenta and the contralateral uterus extending around the fetus, opposite to the placenta. The decidua was defined as SMA-/CD31+/CK19-cells spatially located between the ipsilateral uterus and placenta. The placenta was defined by the presence of CK19+ cekks, indicating fetal trophoblasts. Finally, the fetus was identified as the isolated cluster of cells located within the amniotic cavity/yolk sac. For all implantation sites included in this study, staining for OROV viral antigen was then quantified within these anatomical regions to determine the absence or presence of vertical transmission.

### Statistics

All graphs were plotted using Prism GraphPad (version 11) and statistical analyses were performed using Prism GraphPad or R (version 4.5.2). Data are expressed as mean or medians with scatter plots showing individual values. Number of biological samples used per experiment (n) and statistical tests used for each experiment are included in figure legends. Survival between OROV-infected dams was analyzed using the Mantel-Cox test. Viral loads (RNA and infectious virus) and OROV area localization were analyzed by Mann-Whitney *U* Test and adjusted for multiple comparisons via the Holm-Šídák method when appropriate. Limit of detections are denoted by dotted lines. Fetal body area was analyzed via a Student’s unpaired test. Categorical fetal pathology (intact or resorption) or categorical presence of OROV (positive or negative) were analyzed using a Fisher’s exact test and P values were adjusted for multiple comparisons using the Bonferroni correction when appropriate. p < 0.05 was considered as significant. ns = non-significant, *p < 0.05, **p < 0.01, ***p < 0.001, ****p < 0.0001.

## ACKNOWLEDGEMENTS

We would like to thank Dr. Aaron Ring (Fred Hutchinson Cancer Center) for the gift of recombinant Fc-conjugated mouse IL-27. This work was in part supported by the National Institutes of Health (1R01AI180138-01 to KAJ). Additional funding provided by the Pew Charitable Trusts Biomedical Scholars Program (KAJ), Penn Vet’s Institute for Infectious & Zoonotic Diseases (DTP), the University of Pennsylvania Office of the President (KAJ).

---

**Supplemental Figure 1. Representative images of fetal pathology upon E18.5 after OROV infection.** Representative photos of litters (placenta and matched fetus) on E18.5 from early gestation (blue) dams or mid-gestation dams (green) infected with OROV and either pretreated with isotype control or αIFNAR1 24 hours prior to the start of infection.

**Supplemental Figure 2. Cross A fetal pathology at 7 dpi and E13.5. A.** Images of the entire litter from each IFNAR1^+/-^ dams crossed with IFNAR1^-/-^ sires (Cross A) that were infected in early gestation with OROV. Images were taken after 7 dpi on E13.5. **B.** Fetal size, measured by crown-rump length multiplied by occipital diameter, of intact fetuses derived from WT x WT and IFNAR1^+/-^ x IFNAR1^-/-^ matings (Cross A) at 7 dpi/E13.5.

