## Supplemental Figures for "Fetal-intrinsic antiviral mechanisms emerge over the course of gestation"

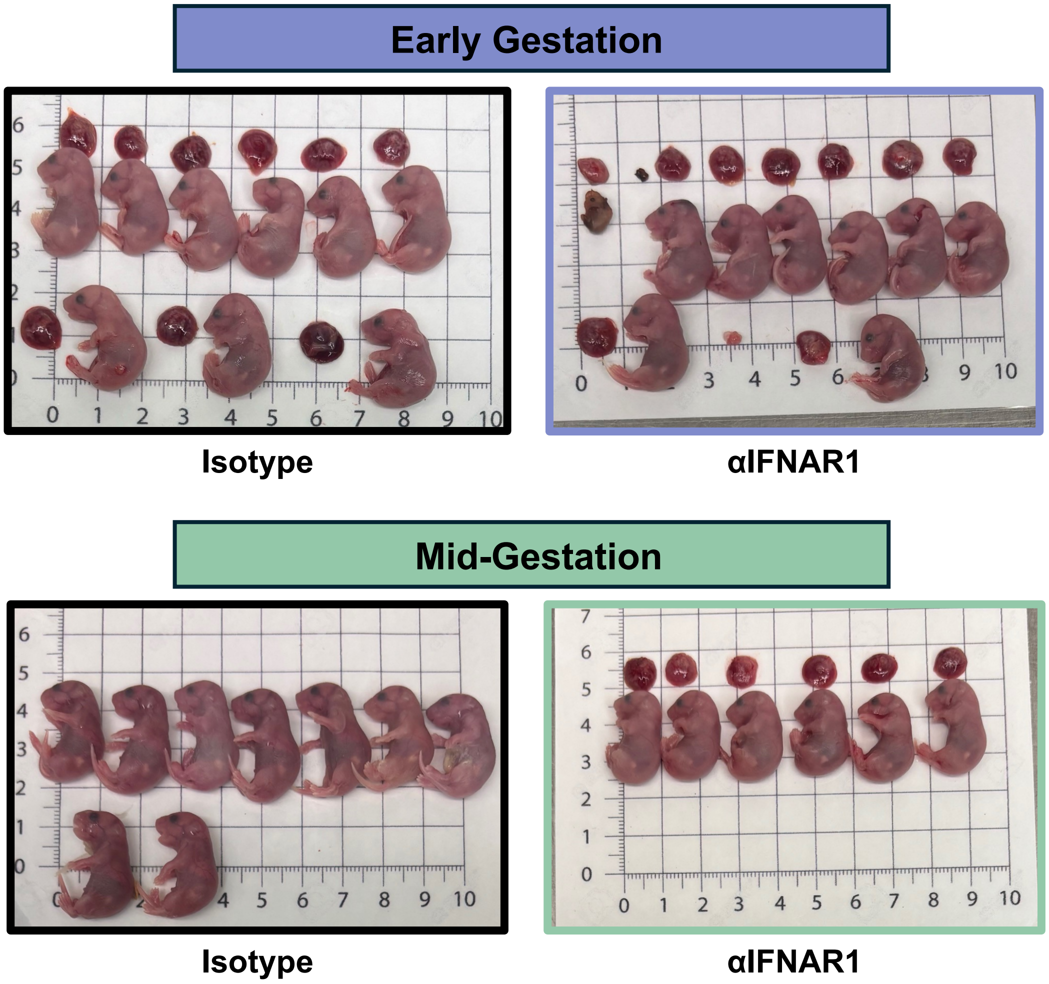


**Supplemental Figure 1. Representative images of fetal pathology upon E18.5 after OROV infection**

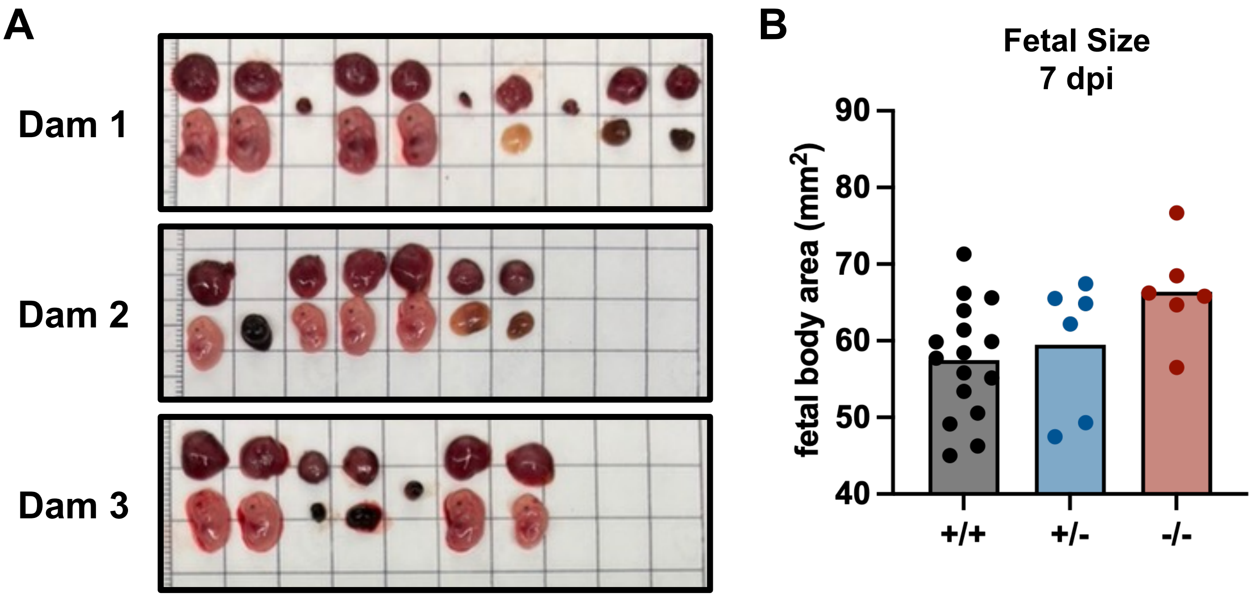


**Supplemental Figure 2. Cross A fetal pathology at 7 dpi and E13.5**
